# OncoContour: An Interactive Platform for Geographic Visualization and Demographic Analysis of Cancer Incidence

**DOI:** 10.64898/2026.05.20.726625

**Authors:** Daniel White, Alper Uzun

## Abstract

Cancer incidence varies substantially across geographic regions and demographic groups, yet translating large-scale surveillance datasets into accessible, interpretable visualizations remains a challenge for researchers and public health professionals without computational expertise. We developed OncoContour, an interactive web-based platform that enables geographic visualization and demographic analysis of cancer incidence data through a browser-accessible interface. To demonstrate its capabilities, we analyzed publicly available cancer incidence data from the United States Cancer Statistics database via CDC WONDER, covering five major cancer types across four northeastern U.S. metropolitan statistical areas from 2017 through 2022, supplemented by demographic data from the U.S. Census Bureau American Community Survey. OncoContour integrates population distribution heatmaps, per-capita cancer incidence heatmaps, interactive multi-city temporal trend charts, structured cancer data tables, and demographic visualizations covering race, ethnicity, age, and sex distributions into a single dynamically generated HTML report. The platform is implemented in Python using Flask, Folium, Plotly, and Matplotlib, and is containerized using Docker for reproducible local deployment. Across all four metropolitan areas, breast and prostate cancers accounted for the highest incidence counts over the study period, while a decline in reported cases observed in 2020 is consistent with documented disruptions to cancer screening during the COVID-19 pandemic. By integrating geospatial mapping, temporal analysis, and demographic visualization within a unified, no-code interface, OncoContour aims to support cancer surveillance, epidemiological investigation, and targeted public health planning. OncoContour is freely available at https://github.com/alperuzun/oncocontour_docker.

## Introduction

Cancer remains one of the leading causes of morbidity and mortality worldwide and continues to impose a substantial burden on health systems and populations. In the United States, cancer is the second leading cause of death overall. Recent estimates indicate that approximately 2,114,850 new cancer cases and 626,140 cancer-related deaths are expected to occur in 2026, corresponding to nearly 5,800 new diagnoses and about 1,700 deaths each day^1^. These numbers highlight the persistent public health impact of cancer despite decades of progress in prevention, early detection, and treatment. Encouragingly, cancer mortality rates have declined substantially since the early 1990s, resulting in an estimated 4.8 million deaths averted through 2023, largely due to reductions in smoking prevalence, improved screening practices, and advances in therapeutic interventions ^1^. Understanding how cancer incidence varies across geographic regions and demographic groups is essential for identifying environmental risk factors, detecting health disparities, and guiding targeted public health interventions. Cancer incidence and mortality rates show substantial variation across geographic regions, reflecting differences in population demographics, environmental exposures, lifestyle factors, healthcare access, and screening practices. For example, incidence rates for preventable cancers such as lung cancer vary several-fold across U.S. states due to differences in smoking prevalence and related risk factors ^1^. Such geographic heterogeneity underscores the importance of spatial analysis in cancer epidemiology and population health research.

Large-scale cancer registries and surveillance programs now generate extensive epidemiological datasets capturing cancer incidence, mortality, and demographic characteristics across populations. Data sources such as the Surveillance, Epidemiology, and End Results (SEER) Program and national cancer registries provide valuable information on cancer occurrence and trends ^2^. However, extracting meaningful insights from these datasets often requires advanced analytical tools and expertise in geographic information systems (GIS), data preprocessing, and statistical analysis. Several widely used platforms support spatial cancer analysis and visualization. For example, the **NCI Cancer Atlas** provides an interactive environment for mapping cancer incidence and mortality across the United States (https://gis.cancer.gov/canceratlas/app/)^3^, while the **CDC United States Cancer Statistics (USCS) visualization tool** enables exploration of cancer trends across regions and demographic groups (https://gis.cdc.gov/Cancer/USCS/) ^4^. In addition, tools such as **SaTScan** (https://www.satscan.org/) support spatial and spatiotemporal cluster detection^5^, and **GeoDa** (https://geodacenter.github.io/) provides advanced spatial data analysis and modeling capabilities widely used in epidemiologic research ^6^.

Many available datasets are distributed in tabular formats that are difficult to interpret without substantial technical expertise, limiting their accessibility to researchers, clinicians, and public health professionals. Geospatial analysis has become an increasingly important component of modern cancer research. Geographic visualization methods enable the identification of disease clusters, regional disparities, and population-level risk patterns. Traditional GIS platforms offer powerful analytical capabilities but often require specialized training, complex software installation, and extensive configuration. Consequently, these tools may be inaccessible to many biomedical researchers and public health practitioners who seek to explore spatial cancer patterns but lack dedicated geospatial expertise. Furthermore, many existing visualization platforms focus primarily on geographic mapping and provide limited integration with demographic, temporal, and comparative analyses. Interactive visualization platforms provide a promising solution by enabling users to dynamically explore complex epidemiological datasets through intuitive graphical interfaces. Integrating geographic mapping with demographic and temporal analyses allows users to examine cancer incidence patterns across multiple dimensions simultaneously, facilitating hypothesis generation and supporting data-driven decision making. Despite these advantages, there remains a need for accessible platforms that integrate spatial cancer mapping with demographic analysis and interactive exploration capabilities.

To address this need, we developed OncoContour, an interactive geographic cancer data visualization platform designed to explore cancer incidence patterns across regions and demographic groups. OncoContour allows users to upload cancer incidence and demographic datasets and generate a range of interactive visualizations, including geographic heatmaps, demographic distributions, temporal trend analyses, and comparative views. By integrating multiple analytical perspectives within a unified interface, the platform enables researchers and public health professionals to explore spatial cancer patterns without requiring specialized programming or geospatial analysis expertise. By simplifying the analysis and visualization of cancer incidence data, OncoContour provides a practical tool for identifying spatial trends, exploring demographic correlations, and supporting population-level cancer research. The platform aims to assist researchers, clinicians, and public health practitioners in deriving insights that may inform cancer surveillance, epidemiological investigations, and targeted public health interventions. In addition to exploratory epidemiological analysis, the platform enables researchers and public health groups to visualize their own geographic cancer datasets, dynamically explore regional patterns, generate presentation-ready visual outputs, and facilitate data sharing and collaborative interpretation across research environments.

## Materials and Methods

### Implementation

OncoContour is a web-based platform designed for the visualization and analysis of geospatial cancer incidence and demographic data. The system integrates data processing, statistical analysis, and interactive visualization into a unified pipeline, enabling users to upload custom datasets and generate dynamic, map-based insights. The platform is designed to run locally on the user’s machine or within a controlled server environment, allowing secure processing of sensitive epidemiological and demographic data without requiring transmission to external servers.

### Programming languages, frameworks, and libraries

OncoContour is implemented as a Python-based web application using the Flask framework, which serves as the backend for handling user requests, data processing, and routing ^7^. The application is deployed as a lightweight server capable of running locally or in a containerized environment. Data manipulation and preprocessing are performed using the Pandas library, enabling efficient handling of tabular datasets including cancer incidence, demographic, and census data ^8^. Numerical operations and transformations are supported by NumPy ^9^. For visualization, OncoContour utilizes a combination of libraries to support both static and interactive graphics. Matplotlib is used to generate static charts such as pie charts and line plots, which are encoded as images and embedded directly into the web interface ^10^. Plotly is used for interactive visualizations, including bar charts, scatter plots, and time-series analyses, providing features such as zooming, panning, and export functionality ^11^. Geospatial visualization is implemented using the Folium library ^12^, which enables the creation of interactive maps and heatmaps rendered in HTML using Leaflet.js with OpenStreetMap as the underlying geographic map layer ^13^. Additional geospatial support is provided through Geopy ^14^ for coordinate handling and location-based computations. The frontend interface is constructed using HTML, CSS, and JavaScript templates rendered through Flask, allowing for dynamic content updates and user interaction without requiring a separate frontend framework.

### System architecture and deployment

OncoContour follows a modular client–server architecture. The Flask backend manages routing, file uploads, data validation, and visualization generation, while the frontend provides an interactive interface for user input and visualization display. Uploaded data files are temporarily stored in a designated directory (“uploads”), where they are validated and processed. The system supports multiple dataset types, including cancer incidence data, city-level geographic data, race/ethnicity distributions, and age/sex demographic data. The application is designed to run locally using Flask’s built-in development server for testing and analysis. For scalable deployment, it can be configured to run within a containerized environment using Docker^15^, ensuring reproducibility and compatibility across operating systems (Figure1).

**Figure 1.**
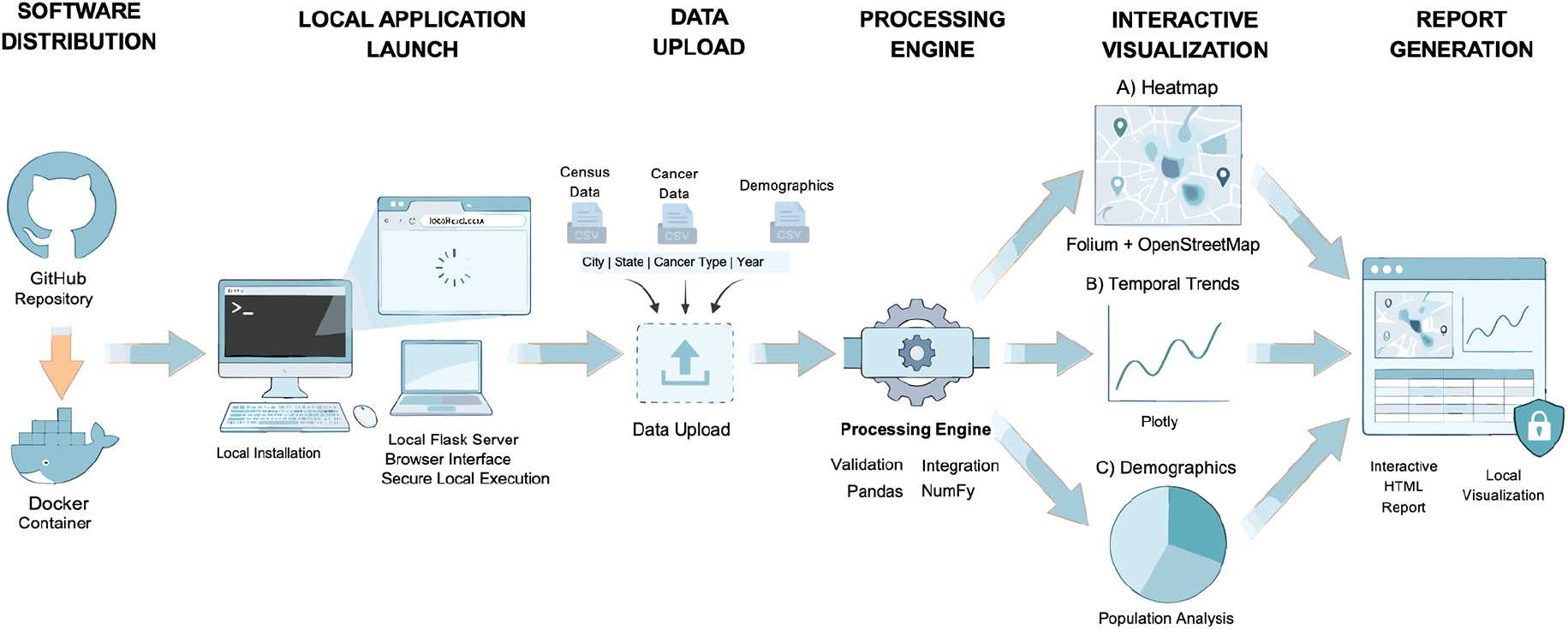
Overview of the OncoContour implementation and deployment workflow. OncoContour is distributed through GitHub as a Docker-compatible application that enables users to install and execute the platform locally on their own machines. After launching the local Flask-based server environment, users upload CSV-based cancer incidence and demographic datasets through a browser-accessible interface. Uploaded data are processed locally using integrated validation, preprocessing, and visualization pipelines implemented with Pandas, Plotly, Folium, and related Python libraries. The platform dynamically generates interactive geospatial heatmaps, temporal trend analyses, demographic visualizations, and unified HTML reports while maintaining secure local data processing without transmission to external servers.

### Data integration and preprocessing

OncoContour integrates user-provided datasets with publicly available census data to enrich geographic and population-level analyses. Specifically, uploaded cancer datasets are merged with processed census data containing city-level population and geographic coordinates. This merging is performed using shared keys such as city and state identifiers, ensuring alignment between epidemiological and demographic data. Input data undergo validation to ensure structural consistency. For example, cancer datasets must include “City” and “State” columns, with state values restricted to valid two-letter U.S. abbreviations. Additional columns are interpreted as either cancer types or yearly incidence values, depending on whether they are textual or numeric. The preprocessing pipeline dynamically identifies cancer-type columns and temporal (year) columns, enabling flexible handling of heterogeneous datasets. Derived metrics, such as total cancer incidence and per-capita cancer rates, are computed during preprocessing to support downstream visualization.

### User input and data ingestion

Users interact with OncoContour through a web-based interface that allows uploading of multiple dataset types. Supported inputs include: **(1)** cancer incidence data by city and year, **(2)** city-level datasets with geographic coordinates, **(3)** county-level race and ethnicity data, and **(4)** age and sex population distributions. Each dataset is uploaded as a CSV file and processed independently. The system allows users to upload one or more datasets simultaneously, enabling flexible analysis workflows. Uploaded files are validated, stored temporarily, and then passed to the visualization pipeline upon user request.

### Visualization and analysis pipeline

The visualization pipeline is initiated upon user request and dynamically generates outputs based on available datasets. The system conditionally constructs visualizations depending on which data files are provided, allowing for modular and adaptive analysis. Geospatial visualizations include population heatmaps and cancer incidence heatmaps. Population heatmaps are generated using latitude, longitude, and population density, while cancer heatmaps display incidence rates normalized per capita. These visualizations enable spatial comparison of disease burden across regions. Interactive charts are generated to analyze demographic and temporal patterns. These include bar charts for race/ethnicity distributions by county, grouped bar charts for age and sex population distributions, line charts for multi-year cancer incidence trends across cities, and pie charts illustrating the distribution of cancer types. Plotly-based visualizations support interactive exploration, while Matplotlib-generated figures provide lightweight embedded summaries. All outputs are compiled into HTML files and rendered within the application interface.

### Dynamic report generation

OncoContour compiles all generated visualizations into a unified HTML report. This report includes geospatial maps, demographic charts, and temporal analyses organized into structured sections. The report is dynamically generated based on available data and can be viewed directly within the browser. The modular design ensures that only relevant visualizations are included, improving usability and clarity. Each visualization is accompanied by contextual information, enabling users to interpret results without requiring additional preprocessing or external tools.

### Secure and local data processing

OncoContour is designed to operate in a local or controlled server environment, ensuring that sensitive health and demographic data remain secure. All uploaded files are processed locally, and no data is transmitted to external servers. This design prioritizes data privacy and compliance with research and clinical data handling standards. Temporary files are managed within the application directory and can be cleared programmatically to prevent data persistence beyond the user session. This approach ensures both security and efficient resource management.

## Results

### Case study: Multi-city cancer incidence analysis (2017–2022)

To evaluate the functionality and analytical capabilities of OncoContour, we conducted a case study using publicly available cancer incidence data from the United States Cancer Statistics (USCS) database via CDC WONDER^16^. Four metropolitan statistical areas (MSAs) in the northeastern United States were selected: the Boston-Cambridge-Newton (MA-NH), Providence-Warwick (RI-MA), Worcester (MA), and Portland-South Portland (ME) metropolitan areas. Five cancer types were analyzed: female breast, prostate, lung and bronchus, colon and rectum, and melanoma of the skin. Data was collected for the years 2017 through 2022. Individual datasets were exported separately from CDC WONDER and transformed into a unified format compatible with OncoContour.

Across all MSAs, breast and prostate cancers accounted for the highest incidence counts over the study period, while melanoma consistently exhibited the lowest counts. The Boston metropolitan area showed the highest overall cancer burden across all five cancer types, reflecting its larger underlying population. Boston exhibited total incidence counts approximately three times higher than Providence and more than six times higher than Portland across the study period. Portland exhibited the lowest total incidence counts, and Providence and Worcester demonstrated intermediate levels with broadly similar distributions across cancer types.

### Geographic visualization of cancer incidence

OncoContour generated two interactive Folium-based heatmaps from the uploaded dataset. The population distribution heatmap rendered intensity proportional to the census-derived population of each city, with Boston appearing as the dominant hotspot. The cancer incidence heatmap normalized total case counts per 100,000 residents, enabling per-capita comparison across regions of different sizes (Figures 2A and 2B). Each map includes interactive markers at each city centroid. Clicking a marker opens a popup displaying a structured table of cancer type counts and population figures for that location (Figure. 2C). The contrast between raw population heatmaps and normalized incidence heatmaps highlights the importance of per-capita adjustment, as regions with smaller populations may exhibit comparable or higher incidence rates despite lower absolute case counts.

**Figure x.**
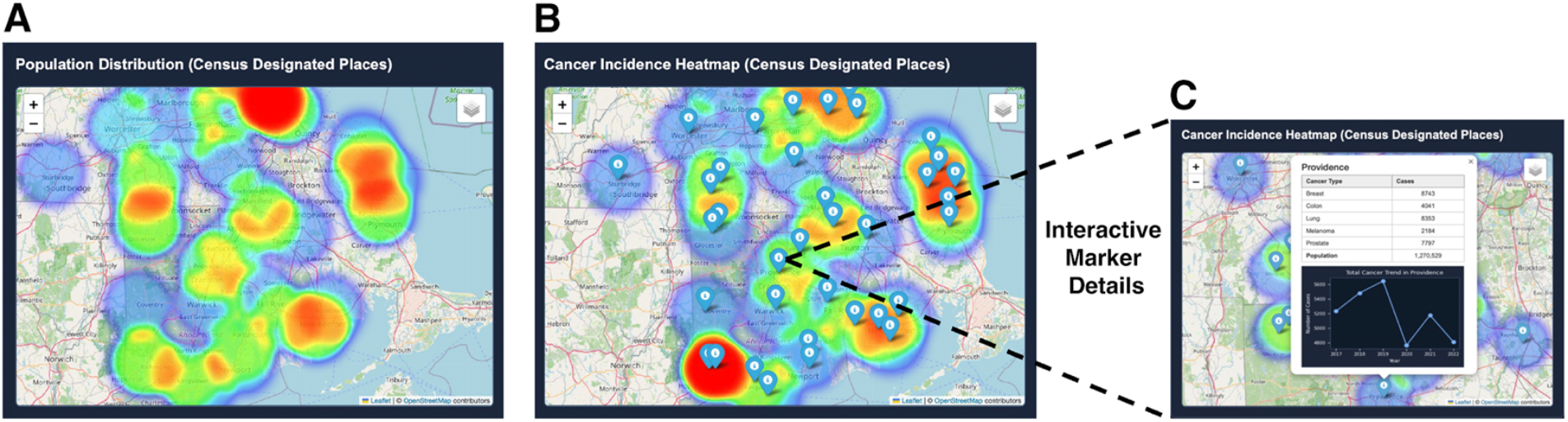
Geographic visualization of population distribution and normalized cancer incidence across census-designated places. **A)** Population distribution heatmap showing spatial variation in population density across the mapped region. **B)** Cancer incidence heatmap normalized per 100,000 individuals and overlaid with interactive location markers representing census-designated places. Warmer colors indicate areas with higher normalized cancer incidence concentration. **C)** Enlarged view of the interactive marker pop-up shown in panel B. Selecting a marker opens a summary window displaying location-specific cancer incidence by cancer type, population statistics, and temporal cancer incidence trends. OpenStreetMap was used as the base map for geographic visualization.

### Temporal trends in cancer incidence

OncoContour produced two types of temporal visualizations from the uploaded data. First, an interactive Plotly line chart compared total annual cancer case counts across all four MSAs from 2017 through 2022. Hover tooltips display the city name, year, and exact case count for each data point. Second, individual Matplotlib line charts were generated for each city and arranged in a two-column grid within the report, providing city-level trend summaries alongside the comparative view (Figure 3). A decline in total reported cases was observed in 2020 across all four MSAs, followed by partial recovery in 2021 and 2022. Boston recorded a peak of 15,155 total cases in 2021 and a decline to 12,740 in 2022. Providence peaked at 5,645 cases in 2019 before declining to 4,767 in 2020. Portland showed more stable trends, increasing modestly from 1,823 cases in 2017 to 2,106 in 2022.

**Figure 3.**
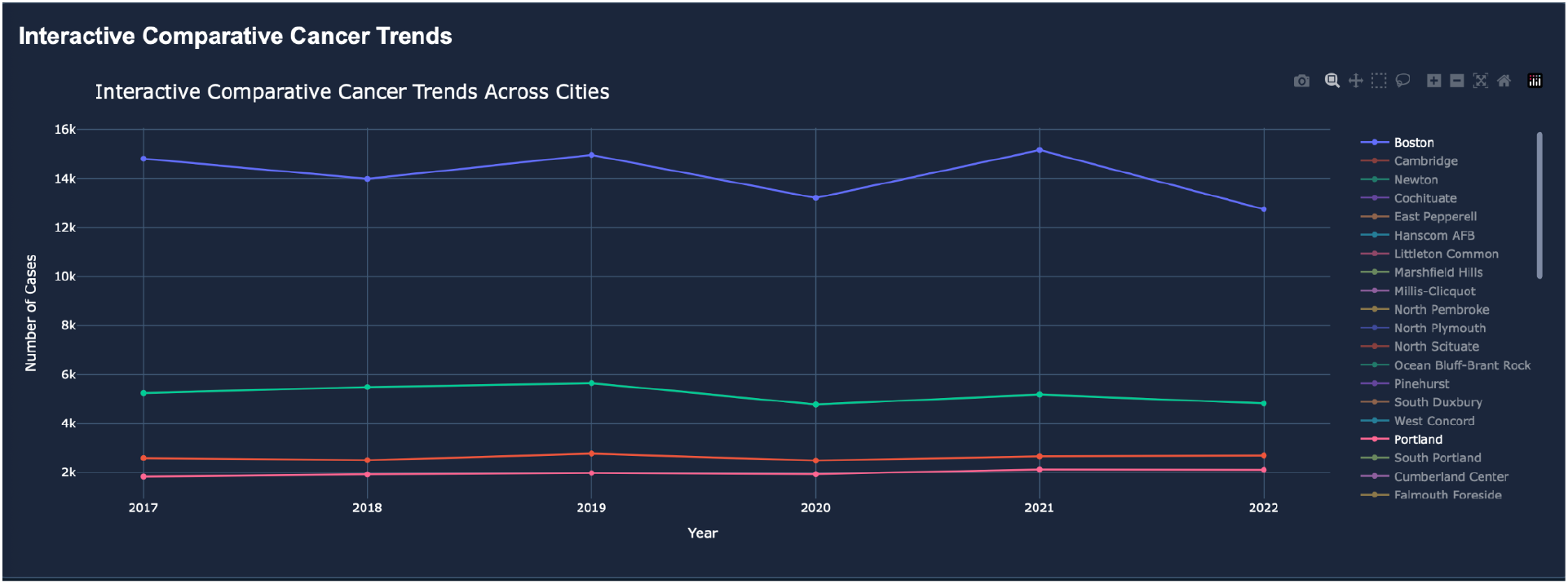
Cancer incidence trend visualizations generated by OncoContour. Interactive Plotly line chart comparing total annual case counts across all four MSAs, with hover tooltips.

### Cancer data table

OncoContour rendered the uploaded cancer incidence data as a structured HTML table within the report, displaying case counts by cancer type for each city. This view provides direct numeric comparison across regions and cancer types. Over the full 2017 to 2022 study period, the Boston MSA recorded 26,081 breast cancer cases, 20,902 prostate cancer cases, 20,238 lung cancer cases, 11,004 colon and rectum cancer cases, and 6,612 melanoma cases. Despite differences in total case volume driven by population size, the relative proportions of cancer types were broadly consistent across all four regions, with breast and prostate cancers consistently representing the largest shares of total incidence.

### Integrated report generation and usability

All visualizations were compiled by OncoContour into a single HTML report rendered within the browser. The report organized outputs into labeled sections: geospatial heatmaps, the cancer data table, the interactive comparative trend chart, and the per-city trend chart grid. The report was generated dynamically based on which data files were uploaded, so only relevant sections were included. No programming knowledge was required beyond uploading the input CSV files through the Import Data page. This design makes the platform accessible to researchers and public health professionals without computational expertise.

### Demographic Distributions and Contextual Analysis

To contextualize the raw spatial and temporal incidence figures, OncoContour successfully processed and visualized localized demographic data for the analyzed MSAs. The racial and ethnic compositions of these regions, highlighting significant demographic variance, were mapped successfully. As shown in Figure 4A, the aggregated racial breakdown across the four MSAs is dominated by White residents (approximately 67.8%), followed by Hispanic or Latino residents (approximately 12.5%), residents identifying as Other (approximately 7.1%), Asian and Pacific Islander residents (approximately 6.7%), and Black residents (approximately 5.9%). At the county level, the Boston-Cambridge-Newton MSA is the most populous region at approximately 4.9 million residents, followed by the Providence-Warwick MSA at approximately 1.6 million, the Worcester MSA at approximately 986,000, and the Portland-South Portland MSA at approximately 563,000 ^17^. The aggregated data confirms that approximately 87.5% of the combined population across these four metropolitan areas identifies as Non-Hispanic or Latino.

**Figure 4.**
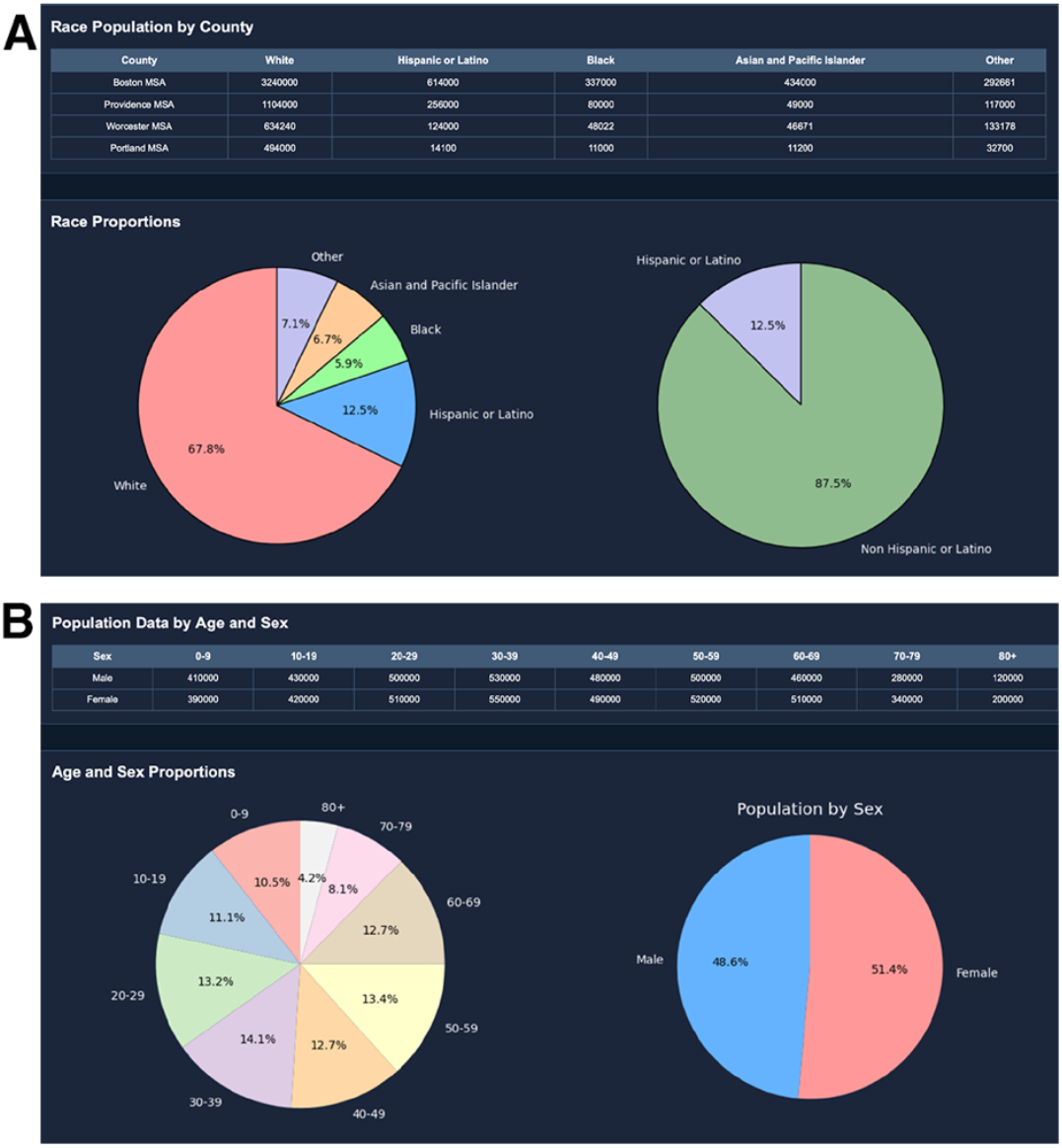
Demographic characterization of analyzed metropolitan statistical areas. **A)** County-level population summary table for the Boston-Cambridge-Newton, Providence-Warwick, Worcester, and Portland-South Portland metropolitan statistical areas. **B)** Demographic distribution visualizations generated by OncoContour, including aggregated racial composition, Hispanic/Latino versus Non-Hispanic/Latino proportions, age cohort distributions, and male/female sex composition across the analyzed regions.

Furthermore, the application generated pie charts representing age and sex distributions across these regions ^17^. The age distribution reveals the largest population concentrations in the 30–39 (14.1%), 40–49 (13.4%), and 50–59 (12.7%) age brackets, with the 50-to-79 cohorts collectively representing a substantial segment of the regional population (Figure 4B). The sex distribution is nearly balanced, with females comprising approximately 51.4% and males approximately 48.6% of the total population across the four MSAs. Correlating this demographic structure with the spatial cancer data^16^ provides immediately actionable insights. Specifically, the elevated incidence rates of age-associated malignancies, such as prostate and breast cancer, strongly align with the aging population clusters identified across these MSAs. By successfully overlaying raw incidence counts with this localized demographic context, OncoContour demonstrates its utility in enabling public health officials to dynamically identify high-risk population subsets and strategically allocate targeted screening and prevention resources.

## Discussion

OncoContour demonstrates that integrating geospatial mapping, demographic analysis, and temporal trend visualization into a single, browser-accessible platform is both technically feasible and practically valuable for cancer epidemiology research. The case study presented here, spanning five cancer types across four northeastern U.S. metropolitan areas from 2017 to 2022, illustrates how the platform enables users to move fluidly between population-level geographic patterns and granular, city-level incidence trends without requiring programming expertise or dedicated GIS software. The observed 2020 dip in reported cases across all four MSAs is consistent with well-documented disruptions to cancer screening and diagnosis during the COVID-19 pandemic, a phenomenon reported broadly in the literature and reflected in national surveillance data ^18^. This finding underscores the value of temporal visualization tools that can surface such anomalies in longitudinal datasets, prompting further investigation.

That said, several important limitations of the platform and the data it relies on warrant discussion. Perhaps the most significant constraint facing OncoContour, and geospatial cancer analysis tools more broadly, is the limited accessibility of granular, publicly available cancer incidence data. In the United States, the most comprehensive cancer surveillance resources, such as the SEER Program and the CDC’s USCS database, provide data aggregated primarily at the county or metropolitan statistical area level, and fine-grained city-or census-tract-level data are frequently suppressed to protect patient privacy when case counts fall below reporting thresholds^2,16^. This suppression is necessary and appropriate, but it meaningfully constrains the spatial resolution at which analyses like those presented here can be conducted. As a result, OncoContour’s heatmaps and per-capita incidence calculations reflect the population centers of the selected MSAs rather than true neighborhood-level variation in cancer burden, which is where health disparities are often most pronounced. Future work could explore integration with state-level cancer registry data, which in some cases offers finer geographic granularity, as well as linkage with social determinants of health datasets to contextualize incidence patterns within broader socioeconomic frameworks.

A related limitation concerns data currency. Because OncoContour relies on user-uploaded CSV files rather than a live API connection to a surveillance database, the recency of the visualizations is entirely dependent on the data the user provides. The USCS database, for instance, typically lags two to three years behind the current year due to the time required for case ascertainment, registry submission, and quality review ^16^. Establishing direct API-based data ingestion from sources like CDC WONDER or the NCI’s SEER*Stat database would significantly improve both the timeliness and reproducibility of analyses conducted through the platform^2,16^. On the technical side, the current implementation runs as a local Flask development server, which is well-suited for exploratory research but presents limitations for broader deployment. The use of Flask’s built-in server is not recommended for production environments handling concurrent users, as it is single-threaded and lacks the performance and security hardening of production-grade web servers^7^. While OncoContour supports containerized deployment through Docker, a more robust production architecture would likely involve a WSGI server such as Gunicorn paired with a reverse proxy like Nginx, along with persistent database storage to replace the current temporary file management approach. Authentication and role-based access controls would also be necessary before the platform could be deployed in clinical or institutional research settings where data governance requirements apply.

From a visualization standpoint, OncoContour currently supports a defined set of chart types, including heatmaps, line charts, bar charts, and pie charts, which cover the most common epidemiological display needs but leave room for expansion. Spatial cluster detection, for example, is a common analytical goal in cancer epidemiology and is currently supported only indirectly through visual inspection of heatmaps. Integrating statistical cluster analysis methods, such as those implemented in SaTScan ^5^ or spatial autocorrelation measures like Moran’s I, would substantially enhance the platform’s analytical depth. Similarly, the current demographic analysis relies on separately uploaded census files derived from American Community Survey estimates ^17^; tighter integration with census APIs could automate demographic enrichment and reduce the burden on users to source and format ancillary data themselves. Support for county-level Federal Information Processing Standards (FIPS) code matching, rather than relying solely on city and state string matching, would also improve the robustness of data merging, particularly for datasets that use inconsistent place-name conventions.

It is also worth noting that the platform’s current case study focuses exclusively on the northeastern United States. While this geographic scope was appropriate for a proof-of-concept evaluation, it does not capture the full diversity of cancer incidence patterns across the country, where regional variation in risk factors such as tobacco use, obesity prevalence, ultraviolet radiation exposure, and access to screening programs drives substantial heterogeneity in incidence and mortality^1^. Expanding the platform’s tested scope to include rural regions, the South, and the Mountain West, areas that often exhibit elevated rates of preventable cancers and face persistent screening access challenges, would be an important direction for future work.

Finally, while OncoContour is designed to be accessible to users without computational backgrounds, the requirement to manually format and upload multiple CSV files with specific column naming conventions represents a non-trivial barrier for non-technical users. A guided data import workflow with validation feedback, pre-loaded example datasets, and support for direct export from common surveillance platforms would meaningfully lower this barrier and broaden the platform’s potential user base.

## Conclusion

OncoContour provides an accessible, browser-based platform for the integrated geographic and demographic visualization of cancer incidence data. By combining Folium-based geospatial heatmaps, interactive Plotly trend charts, and demographic pie chart analyses within a unified Flask application, the platform enables researchers and public health professionals to explore multi-dimensional cancer surveillance datasets without requiring expertise in GIS software or programming. The case study presented here, using CDC WONDER cancer incidence data for four northeastern metropolitan statistical areas, demonstrates the platform’s ability to surface temporal anomalies, support per-capita geographic comparisons, and contextualize incidence patterns within local demographic structures. Limitations related to data granularity, deployment scalability, and the restricted public availability of fine-grained cancer registry data represent important areas for continued development. Future iterations of OncoContour could benefit from direct integration with surveillance data APIs, expanded statistical analysis features, and a more streamlined data import experience. As cancer surveillance datasets continue to grow in scope and complexity, tools that make them interpretable and actionable for a broad range of users will play an increasingly important role in supporting population-level cancer research and public health planning.

